# Social decision making is influenced by size of shoal but not personality or familiarity in Deccan Mahseer (*Tor khudree*)

**DOI:** 10.1101/2020.04.13.040154

**Authors:** Vishwanath Varma, Abhishek Singh, Jintu Vijayan, VV Binoy

## Abstract

Shoals formed by many piscine species are fission-fusion societies where decisions to leave or join a group can have consequences on the fitness of individuals. Some important factors that determine shoal choice are shoal size, familiarity and species composition. However, individuals and species often exhibit distinct shoaling preferences. Individual differences in shoaling preferences may also be related to personality traits such as boldness and sociability. In this study, we examined the link between shoaling decisions and personality traits in a hatchery reared population of an endangered megafish, the Deccan Mahseer (*Tor khudree*). We found that this fish exhibits a distinct preference for larger shoals at ratios of 1:2 or greater. However, they did not prefer to associate with an isolated familiar individual over unfamiliar ones or with a member of their own species over an invasive species. Moreover, shoaling preferences in individuals did not correlate with their boldness or sociability. These results suggest that hatchery reared mahseers which are reintroduced into natural habitats may shoal with invasive species, negatively affecting their viability. Modifying social behaviour of mahseers by amending rearing practices may be a useful strategy to improve outcomes of restocking interventions.

## Introduction

Shoaling in fishes is strongly linked to foraging efficiency and predator avoidance (Ward et al., 2019; Paijmans et al., 2019; Ioannou et al., 2017). Shoals are fission-fusion societies where individuals switch between groups based on innate preferences or the costs and benefits associated with grouping in different contexts. While shoal size, species composition, body size, familiarity and competitive ability of shoal mates are common criteria used for choosing shoals (Marlin et al., 2019; Seguin and Gerlai, 2017; Kleinhappel et al., 2016; Bartolini et al., 2016), the importance of these factors may vary across individuals or species (Paijmans et al., 2019; Vila Pouca and Brown, 2019). Shoaling preferences exerted by individuals can impact the structure and dynamics of social networks, transmission of information and parasites, foraging success, vulnerability to angling, and ultimately, fitness of the populations (Stephenson, 2019; Camacho-Cervantes et al., 2019; Thambithurai et al., 2018; Hasenjager and Dugatkin, 2017; Atton et al., 2014). Hence, the shoaling behaviour of a species can have ramifications on resilience of its populations to natural and anthropogenic environmental disturbances (Maury, 2017; Croft et al., 2003).

Some studies demonstrate individual variation in shoaling preferences within a population (Hellstrom et al., 2016; Cote et al., 2012) and such divergence may correlate with personality traits of individuals (Lucon-Xiccato and Dadda, 2017; Harcourt et al., 2009). Additionally, linkage between personality and social decision making may influence the ability of individuals to modify their decisions in social contexts since personality is associated with behavioural flexibility (Dubois, 2019; Stamps and Biro, 2018). Among the determinants of shoaling decisions that are known to be associated with personality traits are familiarity, species composition and shoal size (Lucon-Xiccato and Dadda, 2017; Benhaim et al., 2016; Cote et al., 2012). For instance, shy individuals of European sea bass exhibit stronger preferences to shoal with familiar individuals (Benhaim et al., 2016). Similarly, in guppies, less sociable individuals tend to associate more with larger shoals (Lucon-Xiccato and Dadda, 2017). Although social decision making in fishes has been a topic of wider investigation, systematic analyses of how individuals varying in their personality modulate their decision making in different social situations are scarce. Since differences in personality traits could influence the structure and dynamics of shoals, understanding the links between major personality traits and shoaling preferences could help improve protocols for hatchery rearing of restocked populations and management of fisheries resources in natural and manmade aquatic ecosystems (Castanheira et al., 2017; Koolhaas and Van Reenen, 2016).

Deccan Mahseers (*Tor khudree*) are large cyprinids indigenous to the rivers of South and Central India (Pinder et al., 2019). Populations of these fish are in decline due to various natural and anthropogenic changes and are being restocked with thousands of hatchery reared juveniles every year (Ogale, 2002). Hatchery reared individuals of several species typically exhibit deficits in social behaviours which affect their growth and survival post release into the natural habitats (Kydd and Brown, 2009; Chapman et al., 2008). However, no study has examined the social behaviour of mahseers and the first report on the personality traits of hatchery reared populations has recently been published (Varma et al. 2020a). Further threat to their natural populations arises from the presence of alien invasive species such as Tilapia (*Oreochromis mossambicus*) with which they may interact and compete for biologically important resources (Raghavan et al., 2011; Sreekantha and Ramachandra, 2005). Tilapia is regarded as one of the most invasive species with the potential to affect behavioural profiles of sympatric species including native fishes (Binoy et al. 2019; Barki and Karplus, 2016). Shoaling with alien invasive heterospecifics is known to be dependent on personality traits (Way et al., 2015) and may be detrimental to native species (Camacho-cervantes et al., 2017). Hence, there is a growing need for understanding the nature of linkage between personality traits and social behaviour in mahseers.

In this study, we addressed three questions:

1. Are shoaling decisions of hatchery reared Deccan Mahseers influenced by the size of stimulus shoals?
2. Does familiarity with conspecific influence the social decision making in situations involving individual conspecifics and alien invasive heterospecific tilapia?
3. Are individuals consistently different from one another in the decisions made across social contexts and does individual variation in shoal choice depend on their personality trait boldness?

## Methods

### Test animals and husbandry

Two months old fingerlings of Deccan Mahseer were procured from the Karnataka Fisheries Department hatchery, Kodagu, Karnataka State, India. They were maintained as groups of 8 (in aquaria of size 40 • 25 • 27 cm) for 8 months in the laboratory and were 5.5 ± 0.13 cm long (Standard Length; SL ± SD; Varma et al., 2020b) when subjected to experimentation. The heterospecifics used in this study were Mozambique tilapia (*Oreochromis mossambicus*), an alien invasive species with size similar to the subject mahseers, purchased from a professional aquarium keeper in Bangalore. Subject mahseers were not exposed to tilapia prior to the experiments. Three sides of all home tanks were covered with black paper to visually isolate the groups from one another. Steel grids wrapped with mosquito nets were placed over the tanks to prevent the fish from jumping out. Water level of 20 cm and temperature of 25 ± 1°C was maintained in the tanks along with a naturalistic light regime of 12:12 hours (Light-Dark, L:D cycle). The tanks were aerated throughout the day to maintain high levels of oxygen and the water was changed every 10 days. Heterospecific tilapia were also maintained in a similar manner. Both tilapia and mahseers were fed *ad libitum* with commercial food pellets every day in the evening.

### Measuring boldness

The personality trait boldness of the subject mahseers was quantified using swim-way apparatus as described in Varma et al. (2020a). An aquarium (80 • 40 • 33 cm) was divided into two unequal chambers, A (15 • 40 • 33 cm) and B (65 • 40 • 33 cm) by an opaque plexiglass sheet; the former functioned as the start chamber while chamber B was the open area available for exploration. The partition wall was fitted with a guillotine door (8 • 5 cm). The apparatus was covered with black paper on all four sides while the tank floor was kept white to provide greater visibility of the subject fish in the video recorded for analysis. The water level was maintained at 15 cm and a compact fluorescent lamp (40 W) illuminated the experimental arena from above. The apparatus was cordoned off using black curtains to avoid external disturbances and a video camera fitted above was used for recording the experiments. Boldness was tested by introducing subject fish into chamber A individually. The guillotine door was raised after 5 minutes given for acclimation. The time taken to enter chamber B (emergence latency - a measure of boldness; Toms et al. 2010) was recorded. Fish were given 10 minutes for entering and exploring chamber B and those that did not emerge from chamber A for over 10 minutes were given ceiling values of 600 seconds and the trial was terminated. We tested the boldness of 63 individual fish in this manner.

### Testing social decision making

#### General protocol

We recorded social decision making by the subject mahseers that had already been tested for boldness. The binary choice apparatus used for this experiment was composed of an aquarium of dimensions 80 • 40 • 33 cm split into three chambers using a transparent plexiglass partition (Binoy et al. 2015). The side chambers (15 • 40 • 33 cm) were used for presenting the stimulus shoals while the subject fish was introduced in the central chamber (50 • 40 • 33 cm). A transparent presentation cage made of plexiglass (15 • 11 • 33 cm) was used for confining the subject fish in the middle of the central area to enable visual observation of the stimulus shoals present on both sides prior to the release. This whole apparatus was also covered on all four sides with black paper and isolated from the rest of the room by black curtains to minimize disturbances.

The stimulus shoals were placed in the side chambers of the binary choice apparatus and allowed to acclimatize for 5 minutes. The subject fish was then introduced into the presentation cage kept in the central chamber and 5 minutes were given for the assessment of the stimulus shoals. Subsequently, the holding chamber was raised and the subject fish was allowed to choose between the available stimuli shoals for a duration of 10 minutes. The time spent within 5 cm of either of the two plexiglass partitions separating side chambers from the central arena of the apparatus (preference zone) was recorded. All stimulus fish were selected carefully to ensure that they were of similar size as the subject fish to control for size dependent shoaling preference (Webster et al., 2007).

#### Influence of shoal size

The influence of shoal size on social decision making by mahseer was tested by providing the following combinations of different numbers of unfamiliar conspecifics in the two side chambers: 2 vs. 0 (ratio 1:0; here one side chamber was kept empty to estimate sociability), 2 vs. 2 (1:1), 2 vs. 3 (1:1.5), 2 vs. 4 (1:2), and 2 vs. 8 (1:4). Individual fishes (N=34) were tested following the protocol mentioned above and the sequence of the trials was randomized.

### Influence of species and familiarity with conspecific

Three types of stimulus fish i.e. familiar conspecifics, unfamiliar conspecifics and unfamiliar heterospecifics were used in this experiment. Three different combinations were tested by providing individual fishes in the side chambers; familiar mahseer vs unfamiliar mahseer, unfamiliar mahseer vs unfamiliar tilapia, or familiar mahseer vs unfamiliar tilapia. We used a separate set of subject fish (N=29) for this experiment. Akin to the group choice experiment, boldness of these fish was also quantified before the binary choice assay. Familiar conspecifics were fish that had cohabited with the focal fish in their home tank for 8 months whereas unfamiliar conspecifics or heterospecifics are fish which have not previously encountered the focal fish. Since no previous information on the number of days required to develop familiarity is available for mahseer, this duration of cohabitation was provided to ensure the acquisition of familiarity.

### Statistical Analysis

All behaviours were quantified from the video recording using BORIS v.6.2.2 (Behavioral Observation Research Interactive Software; Friard and Gamba, 2016). The Preference Index (PI) was calculated for the shoal choice experiments by following the formula from Gomez-Laplaza and Gerlai, 2012: PI = (Time spent along larger shoal) / (Total time spent along both stimulus shoals). In experiments where individual fish were used as stimuli, time spent near the familiar individual was used as the numerator and the sum of the total time spent with both fishes formed the denominator in the formula used to calculate PI. Similarly, in the experiment assessing shoaling preference for conspecific vs heterospecific, time spent near the former was used as the numerator. PIs were tested for normality using Shapiro-Wilk’s test and significant difference from chance using one-sample t-test against a value of 0.5. Since the data followed normal distribution, Linear Mixed Models (LMM) were used to analyse preference of shoal size using preference index as response variable, boldness, sociability (PI for fish shoal over empty chamber in 1:0 condition) and treatments (1:1, 1:1.5, 1:2 and 1:4 ratios of shoal sizes) as independent variables, and FishID (identity of individual fish) as random effect. LMM using PI as response variable, boldness and treatments (familiar vs unfamiliar conspecific, unfamiliar conspecific vs unfamiliar heterospecific, familiar conspecific vs unfamiliar heterospecific) as independent variables, and FishID as random effect were used to analyse the preference for familiarity and species. Full models including all possible effects were first analysed before sequentially dropping non-significant effects. Better fits of models without a particular variable with respect to full models (as determined by ΔAIC > 2, where the model with the lower AIC is considered a better fit) were taken as evidence of the significance of the dropped variable. Residuals after fitting the model were verified to be normally distributed by visual inspection. *Post-hoc* comparisons using Tukey’s Honest Significant Difference test were performed to compare multiple treatments. All statistical analysis was performed using ‘lsmeans’ and ‘lmer’ packages on R.

## Results

### Shoal Size Preference

Linear mixed models using Preference Index as response variable revealed a positive slope for increase in ratio of shoal size (β = 0.07 for 1:1.5, β = 0.2 for 1:2 and β = 0.3 for 1:4) with *post-hoc* multiple comparisons using Tukey’s HSD test revealing significantly higher preference for the larger shoal in 1:2 and 1:4 ratios compared to 1:1 (p < 0.001; Fig. 1a). However, there was no significant difference in PI between 1:1 and 1:1.5 ratios (p > 0.05; Figure 1a). When mahseers were allowed to choose between a shoal of 2 individuals and an empty chamber (1:0 ratio), they showed a clear preference for the shoal with a mean PI of 0.71 (p < 0.001, t-test against 0.5). The PI for each individual in this condition was used as the sociability value. The variation in the PI across treatments that was explained by FishID was negligible (Variance = 0.00069) and the model without the random effect of individuals was a better fit than the model with it (ΔAIC = 2; p > 0.05), suggesting that individuals are not consistent in their preference for the larger shoal across different ratios. Moreover, LMMs with boldness (β = −0.00001; Figure 1b) or sociability (β = 0.1) were not better fits than those without (ΔAIC < 2). These results suggest that preference for the larger shoal is apparent only at ratios greater than 1:1.5 and is not related to personality traits in Deccan Mahseers.

**Figure 1.**
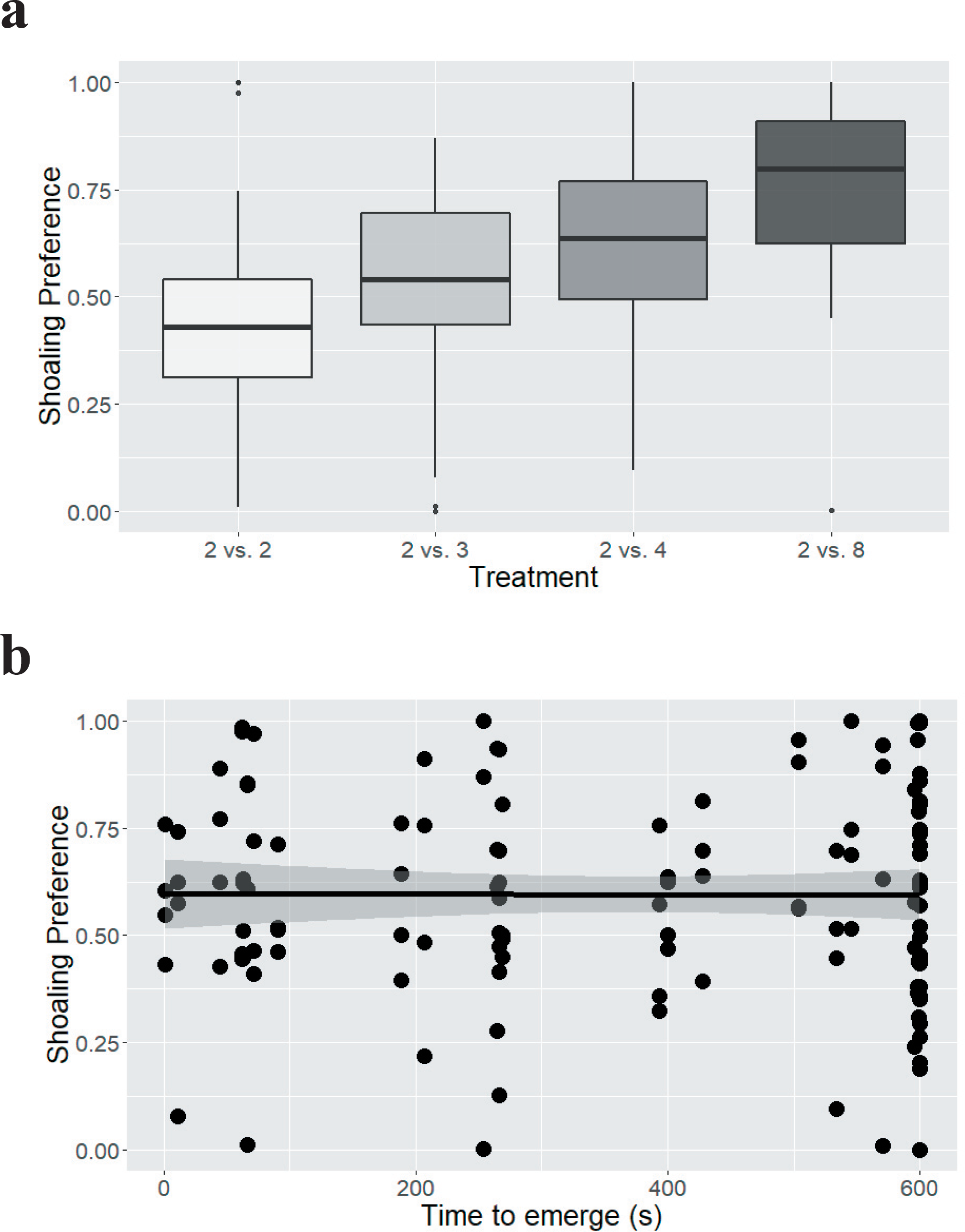
Preference of larger shoal size and its correlation with Boldness. a) Preference index calculated as proportion of time spent near larger shoal with respect to the total time spent shoaling increases with increase in ratio of shoal size. b) Preference index among individuals for larger shoals is not correlated with boldness (time to emerge). Regression line is plotted to indicate relationship between the two variables in panel b.

### Familiarity and Conspecific Preference

PIs from unfamiliar conspecific vs unfamiliar heterospecific (p = 0.09), familiar conspecific vs unfamiliar conspecific (p = 0.8) or familiar conspecific vs unfamiliar heterospecific (p = 0.9) did not yield values significantly different from chance (t-test comparing preference index to 0.5) suggesting lack of discrimination between shoals based on familiarity or species in these fish (Figure 2a). Furthermore, LMM on preference index across familiarity and species revealed no significant effect of boldness (β = 0.00006; p > 0.05; Figure 2b). PI did not vary across different social contexts either, with *post-hoc* comparisons using Tukey’s HSD test revealing no significant differences between the treatments (p > 0.05). No consistency was observed in the preference shown by individual fish for familiar individuals over unfamiliar individuals or the conspecific over the heterospecific since the model without the random effect is a better fit of the dataset (ΔAIC = 2; p > 0.05). Hence, we do not find evidence for shoaling preference based on familiarity or species when stimuli are individuals and these shoaling decisions does not appear to be consistent among individual fish or related to their boldness.

**Figure 2.**
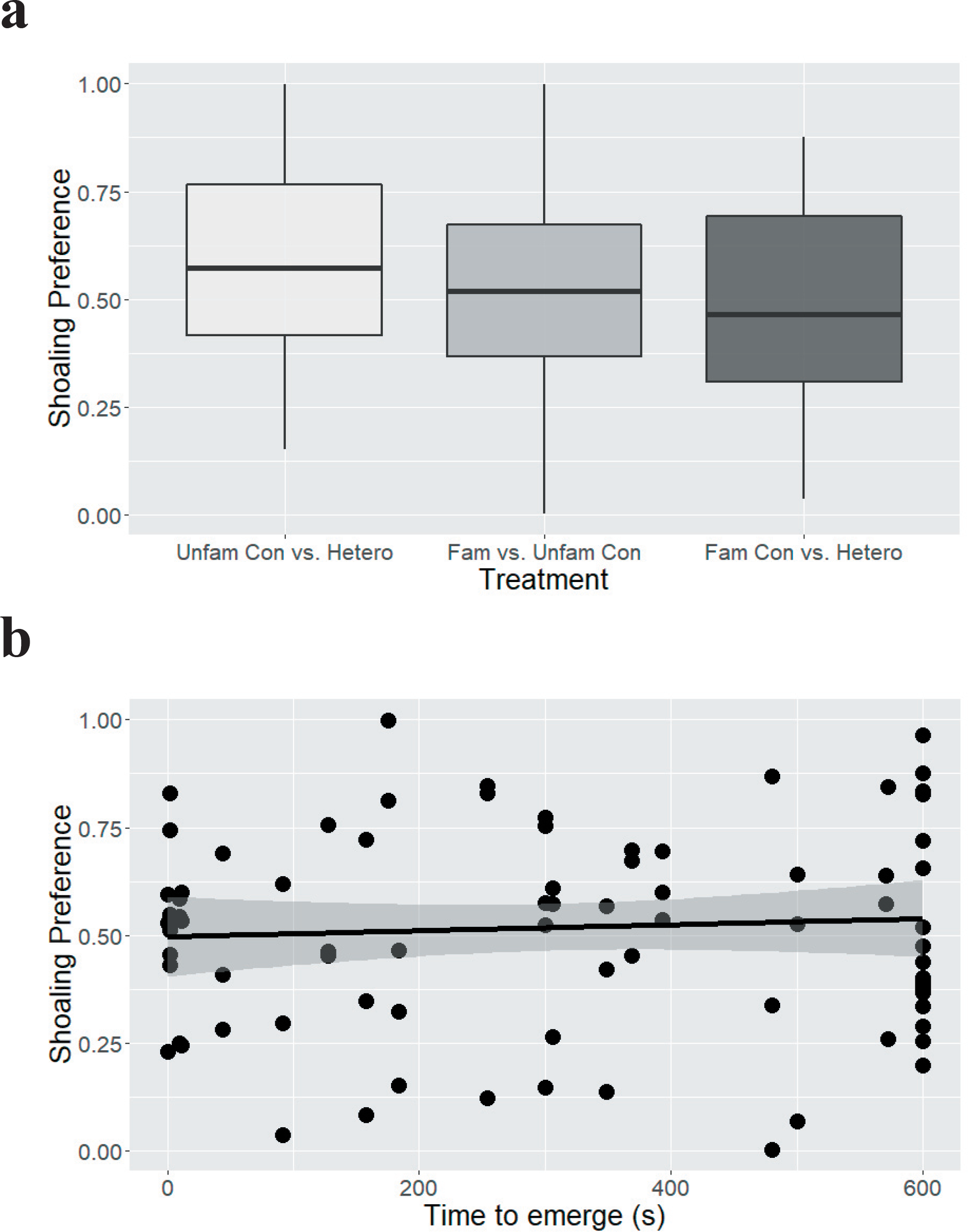
Preference of Species or Familiarity and its correlation with Boldness. a) Preference indices for unfamiliar conspecifics (UC) over unfamiliar heterospecifics (UH), for familiar conspecifics (FC) over unfamiliar conspecifics, and for familiar conspecifics over unfamiliar heterospecifics are not significantly different from chance (0.5) suggesting lack of preference based on familiarity or species. b) Preference index calculated as proportion of time spent near familiar or unfamiliar conspecific with respect to the total time spent shoaling is not correlated with time taken to emerge into a novel environment (boldness, in seconds).

## Discussion

Hatchery reared juveniles of Deccan Mahseers preferred to join shoals with greater number of conspecifics similar to previous reports in many other piscine species (Binoy and Thomas 2004; Pritchard et al., 2001; Krause and Godin, 1994). Being the member of a large group provides benefits of collective vigilance and reduced predation risk (dilution effect and many eyes hypothesis; Rodgers et al., 2013; Boland et al., 2003) and group foraging (Krause et al., 2002) though they have to encounter greater competition and increased danger of parasitic and pathogenic infections (Grand & Dill, 1999). Mahseer juveniles are vulnerable to predation with reports indicating the sympatric occurrence of various predatory species (Prasad et al., 2009). Although shoaling behaviour of the juveniles of Deccan Mahseer in the natural habitat is not well studied, large aggregations of adults and subadults of this species have been seen at several sites (Kumar and Devi, 2013). Interestingly, mahseers which were unable to reach a consistent shoaling decision when the ratio of the stimulus shoal were 1:1 and 1:1.5, expressed a significant preference towards the larger shoals when the ratio of shoal size was 1:2 or greater. The ability to distinguish between numerical values is referred to as numerical acuity and is known to occur in several animals including fishes (Agrillo et al., 2017). However, the numerical acuity of juvenile mahseers from this study is marginally lower (1:2) than guppies and sticklebacks which clearly prefer the larger shoal at ratios of 1:1.25 and 1:1.16 respectively (Agrillo et al., 2017). However, further studies with similar ratios but different absolute numbers of individuals constituting the shoal are required to obtain a clearer estimate of the numerical ability in this species.

Preference to join conspecifics over heterospecifics is widely observed across fish species due to the benefits of phenotypic homogeneity which results in reduced likelihood of predation (oddity effect; Aivaz et al., 2019; Landeau and Terbogh, 1986). However, some species such as sticklebacks and killifish do not always show preference for conspecifics over heterospecifics, depending on shoal size, diet similarity or the extent of phenotypic similarity between the two species (Kleinhappel et al., 2016; Krause and Godin, 1994). Although mahseers preferred larger groups of conspecifics to shoal with, they failed to exhibit a preference for an isolated member of its own species when presented in combination with an alien invasive heterospecific tilapia. In our protocol, we ensured similarity in all factors (such as diet and body size) other than species itself that may influence shoaling decisions in order to ascertain whether individuals choose their shoaling partners based on species alone. In this context, mahseers did not show preference for conspecifics over tilapia. The inability to discriminate conspecific from heterospecific may be attributed to the minimal differences in visual features of Deccan Mahseers and tilapia. Alternatively, the lack of species specific shoaling in Deccan Mahseers could also be due to possible benefits of cross-species sociability (Sridhar and Guttal, 2018).

Previous experience with conspecifics and heterospecifics can also bias shoaling decisions such that fishes often spend more time near familiar individuals over unfamiliar ones (Griffiths et al., 2003). However, the subject species mahseer failed to exhibit a significant preference towards either stimulus when unfamiliar tilapia was presented in combination with an individual conspecific which had cohabited with the subject fish for 8 months. Moreover, no preference towards the familiar conspecific over unfamiliar conspecifics was observed in Deccan Mahseer. Although the time required for acquiring familiarity may vary from species to species, it is unlikely that the number of days the individuals have spent together in our study is insufficient to develop familiarity (Jacoby et al., 2012; Binoy and Thomas, 2004). Instead, the birth and maintenance of the subjects of the present study in captive conditions might have negatively impacted their social decision making. This is consistent with a previous report that preference for familiar individuals was lost in captive and laboratory reared populations of rainbowfish (Kydd and Brown, 2009). The presence of minimal environmental complexity and the absence of selection pressure in the hatchery conditions may be responsible for such behavioural and cognitive deficits in fish (Salvanes et al., 2013). Alternatively, Deccan Mahseer as a species may be incapable of acquiring familiarity with conspecifics and heterospecifics. This pertains to the trade-off between the benefits of recognizing and interacting with familiar individuals in reducing aggression and competition (Hojesjo et al., 1998) and the physiological costs of maintaining cognitive machinery required for individual recognition in fishes (Griffiths, 2003). If Deccan Mahseers live in unstable groups with high rates of fission and fusion among shoals, the benefits of individual recognition would be outweighed by its costs. While high predation environments tend to create stable and strong social networks in guppies (Kelley et al., 2011), there is no study examining stability of social groups in Deccan Mahseers. Therefore, comparison of hatchery reared populations with wild populations and observations of social network structure of these fish in their natural habitat would be needed to validate these explanations.

The inability to develop familiarity with conspecifics or prefer conspecifics over alien invasive heterospecifics raises concerns for restocking of this species. Since mahseers share their habitat with alien invasive species such as tilapia (Raghavan et al., 2011) and preferences exhibited in the laboratory are indications of the social structure of shoals in the wild (Krause et al., 2000), restocked mahseers may be expected to attempt to join shoals with unfamiliar conspecifics and heterospecifics. Invasive species are known to compromise the fitness of native species by shoaling with them (Camacho-Cervantes et al., 2019). Further studies will be required to examine whether fitness of mahseers may be negatively impacted by invasive tilapia in the natural habitats and how such effects may be prevented by preparing hatchery reared individuals for situations they have not previously experienced

We also explored the relationship between the personality traits, boldness and sociability, of subject fish with their shoaling decision making. In contrast to some previous studies on fish species such as sticklebacks *Gasterosteus aculeatus* and guppies *Poecilia reticulata* (Barber et al., 2017), boldness was not correlated with decision making in Deccan Mahseers. Similar to a previous study in *Poecilia reticulata* where sociability (or tendency to join a shoal) did not predict number discrimination (Petrazzini and Agrillo, 2016), we also found that in mahseers, shoaling preference was not correlated with sociability. This is consistent with the lack of correlation between personality and lateralisation, previously observed in this species (Varma et al., 2020a). Instead, shoaling decisions may be more strongly linked with motivation rather than personality itself (Lucon-Xiccato and Bisazza, 2017). Shoaling preference may be a balance between preference for smaller shoals due to motivation to avoid competition and preference for larger shoals due to the motivation to minimize predation pressure (Krause and Godin, 1994; Krause, 1993). Different individuals may prefer larger or smaller shoals based on trade-offs between their tendency to avoid competition or to reduce predation risk, rather than exhibit shoal choice based on their personality. Hence, this may not be expressed as a correlation with personality at the population level.

Overall, we find that hatchery reared Deccan Mahseer do not choose shoaling partners based on species matching or familiarity but do prefer to associate with larger shoals. However, these preferences were not consistent within individuals or correlated with personality traits such as boldness and sociability. These results have implications for the behaviour and survival of hatchery reared fish that are reintroduced into natural environments since they may shoal with invasive heterospecifics that could out-compete them. A comprehensive comparison of hatchery reared fish with fish from the natural habitat will be required to determine whether the shoaling phenotype of captive reared fish is caused by the rearing environment. Accordingly, negative consequences of such rearing conditions may be attenuated by adding complexity and mimicking natural environments such as has been demonstrated for exploratory and anti-predator behaviour in golden mahseers (Ullah et al., 2017). Additionally, hatchery reared fish may be trained to exhibit appropriate shoaling behaviours to maximize fitness prior to release into the wild (Chapman et al., 2008; Salvanes et al., 2007). This study provides a primer for subsequent efforts towards replenishing natural populations of Deccan Mahseers with hatchery reared individuals.

## Acknowledgements

The authors are grateful to the Karnataka State Fisheries Department for kindly providing the mahseer fingerlings, and would like to thank Darshan Pramod (District Fisheries Department, Kodagu) and Dr. Rajeev Raghavan for their support. VV thanks the Department of Biotechnology, Government of India, for a Research Associate Fellowship.

## References

Agrillo, C., Petrazzini, M. E. M., & Bisazza, A. (2017). Numerical abilities in fish: a methodological review. Behavioural processes, 141, 161–171.

Aivaz, A. N., Manica, A., Neuhaus, P., & Ruckstuhl, K. E. (2020). Picky predators and odd prey: colour and size matter in predator choice and zebrafish’s vulnerability–a refinement of the oddity effect. Ethology Ecology & Evolution, 32(2), 135–147.

Atton, N., Galef, B. J., Hoppitt, W., Webster, M. M., & Laland, K. N. (2014). Familiarity affects social network structure and discovery of prey patch locations in foraging stickleback shoals. Proceedings of the Royal Society B: Biological Sciences, 281(1789), 20140579.

Barber, I., Mora, A. B., Payne, E. M., Weinersmith, K. L., & Sih, A. (2017). Parasitism, personality and cognition in fish. Behavioural processes, 141, 205–219.

Barki, A., & Karplus, I. (2016). The behavioral mechanism of competition for food between tilapia (Oreochromis hybrid) and crayfish (Cherax quadricarinatus). Aquaculture, 450, 162–167.

Barnard, C. J., & Burk, T. (1979). Dominance hierarchies and the evolution of “individual recognition”. Journal of theoretical Biology, 81(1), 65–73.

Bartolini, T., Mwaffo, V., Showler, A., Macrì, S., Butail, S., & Porfiri, M. (2016). Zebrafish response to 3D printed shoals of conspecifics: the effect of body size. Bioinspiration & biomimetics, 11(2), 026003.

Benhaïm, D., Ferrari, S., Chatain, B., & Bégout, M. L. (2016). The shy prefer familiar congeners. Behavioural processes, 126, 113–120.

Bergmüller, R., & Taborsky, M. (2010). Animal personality due to social niche specialisation. Trends in Ecology & Evolution, 25(9), 504–511.

Binoy, V. V. (2015). Comparative analysis of boldness in four species of freshwater teleosts. Indian J. Fish, 62(1), 128–130.

Boland, C. R. (2003). An experimental test of predator detection rates using groups of freeliving emus. Ethology, 109(3), 209–222.

Bower, S. D., Mahesh, N., Raghavan, R., Danylchuk, A. J., & Cooke, S. J. (2019). Sub-lethal responses of mahseer (Tor khudree) to catch-and-release recreational angling. Fisheries research, 211, 231–237.

Camacho-Cervantes, M., Palomera-Hernadez, V., & García, C. M. (2019). Foraging behaviour of a native topminnow when shoaling with invaders. Aquatic Invasions, 1(3), 140101.

Carere, C., & Locurto, C. (2011). Interaction between animal personality and animal cognition. Current Zoology, 57(4), 491–498.

Castanheira, M. F., Conceição, L. E., Millot, S., Rey, S., Bégout, M. L., DamsgAard, B., … & Martins, C. I. (2017). Coping styles in farmed fish: consequences for aquaculture. Reviews in Aquaculture, 9(1), 23–41.

Chapman, B. B., Ward, A. J., & Krause, J. (2008). Schooling and learning: early social environment predicts social learning ability in the guppy, *Poecilia reticulata*. Animal Behaviour, 76(3), 923–929.

Clarke, G. M. (1998). The genetic basis of developmental stability. V. Inter-and intraindividual character variation. Heredity, 80(5), 562–567.

Cote, J., Fogarty, S., & Sih, A. (2012). Individual sociability and choosiness between shoal types. Animal Behaviour, 83(6), 1469–1476.

Croft, D. P., Krause, J., Couzin, I. D., & Pitcher, T. J. (2003). When fish shoals meet: outcomes for evolution and fisheries. Fish and Fisheries, 4(2), 138–146.

Dubois, F. (2019). Why are some personalities less plastic?. Proceedings of the Royal Society B, 286(1908), 20191323.

Friard, O., & Gamba, M. (2016). BORIS: a free, versatile open[source event[logging software for video/audio coding and live observations. Methods in Ecology and Evolution, 7(11), 1325–1330.

Gómez-Laplaza, L. M., & Gerlai, R. (2012). Activity counts: the effect of swimming activity on quantity discrimination in fish. Frontiers in psychology, 3, 484.

Grand, T. C., & Dill, L. M. (1999). The effect of group size on the foraging behaviour of juvenile coho salmon: reduction of predation risk or increased competition?. Animal Behaviour, 58(2), 443–451.

Griffiths, S. W. (2003). Learned recognition of conspecifics by fishes. Fish and Fisheries, 4(3), 256–268.

Griffiths, S., Höjesjö, J., & Johnsson, J. (2003). Familiarity confers anti[predator and foraging advantages on juvenile brown trout. Journal of Fish Biology, 63, 226–226.

Griffiths, S. W., & Magurran, A. E. (1997). Familiarity in schooling fish: how long does it take to acquire?. Animal Behaviour, 53(5), 945–949.

Harcourt, J. L., Sweetman, G., Johnstone, R. A., & Manica, A. (2009). Personality counts: the effect of boldness on shoal choice in three-spined sticklebacks. Animal Behaviour, 77(6), 1501–1505.

Hasenjager, M. J., & Dugatkin, L. A. (2017). Fear of predation shapes social network structure and the acquisition of foraging information in guppy shoals. Proceedings of the Royal Society B: Biological Sciences, 284(1867), 20172020.

Hellström, G., Heynen, M., Borcherding, J., & Magnhagen, C. (2016). Individual consistency and context dependence in group-size preference of Eurasian perch. Behavioural processes, 133, 6–11.

Herczeg, G., Gonda, A., & Merilä, J. (2009). The social cost of shoaling covaries with predation risk in nine-spined stickleback, *Pungitius pungitius*, populations. Animal Behaviour, 77(3), 575–580.

Hoffmann, A. A., Merilä, J., & Kristensen, T. N. (2016). Heritability and evolvability of fitness and nonfitness traits: lessons from livestock. Evolution, 70(8), 1770–1779.

Höjesjö, J., Johnsson, J. I., Petersson, E., & Järvi, T. (1998). The importance of being familiar: individual recognition and social behavior in sea trout (*Salmo trutta*). Behavioral Ecology, 9(5), 445–451.

Huizinga, M., Ghalambor, C. K., & Reznick, D. N. (2009). The genetic and environmental basis of adaptive differences in shoaling behaviour among populations of Trinidadian guppies, *Poecilia reticulata*. Journal of evolutionary biology, 22(9), 1860–1866.

Ioannou, C. C., Ramnarine, I. W., & Torney, C. J. (2017). High-predation habitats affect the social dynamics of collective exploration in a shoaling fish. Science advances, 3(5), e1602682.

Jacoby, D. M. P., Sims, D. W., & Croft, D. P. (2012). The effect of familiarity on aggregation and social behaviour in juvenile small spotted catsharks Scyliorhinus canicula. Journal of Fish Biology, 81(5), 1596–1610.

Kelley, J. L., Morrell, L. J., Inskip, C., Krause, J., & Croft, D. P. (2011). Predation risk shapes social networks in fission-fusion populations. PloS one, 6(8), e24280.

Kitada, S. (2018). Economic, ecological and genetic impacts of marine stock enhancement and sea ranching: A systematic review. Fish and Fisheries, 19(3), 511–532.

Kleinhappel, T. K., Burman, O. H., John, E. A., Wilkinson, A., & Pike, T. W. (2016). A mechanism mediating inter-individual associations in mixed-species groups. Behavioral Ecology and Sociobiology, 70(5), 755–760.

Koolhaas, J. M., & Van Reenen, C. G. (2016). Animal behavior and well-being symposium: Interaction between coping style/personality, stress, and welfare: Relevance for domestic farm animals. Journal of animal science, 94(6), 2284–2296.

Krause, J., Butlin, R. K., Peuhkuri, N., & Pritchard, V. L. (2000). The social organization of fish shoals: a test of the predictive power of laboratory experiments for the field. Biological Reviews, 75(4), 477–501.

Krause, J., & Godin, J. G. J. (1994). Shoal choice in the banded killifish (Fundulus diaphanus, Teleostei, Cyprinodontidae): effects of predation risk, fish size, species composition and size of shoals. Ethology, 98(2), 128–136.

Krause, J., & Godin, J. G. J. (1995). Predator preferences for attacking particular prey group sizes: consequences for predator hunting success and prey predation risk. Animal Behaviour, 50(2), 465–473.

Krause, J., Ruxton, G. D., Ruxton, G. D., & Ruxton, I. G. (2002). Living in groups. Oxford University Press, UK.

Kumar, R., & Devi, K. R. (2013). Conservation of freshwater habitats and fishes in the W estern G hats of I ndia. International Zoo Yearbook, 47(1), 71–80.

Kydd, E., & Brown, C. (2009). Loss of shoaling preference for familiar individuals in captive•reared crimson spotted rainbowfish *Melanotaenia duboulayi*. Journal of Fish Biology, 74(10), 2187–2195.

Landeau, L., & Terborgh, J. (1986). Oddity and the ‘confusion effect’in predation. Animal Behaviour, 34(5), 1372–1380.

Lucon-Xiccato, T., & Dadda, M. (2017). Personality and cognition: sociability negatively predicts shoal size discrimination performance in guppies. Frontiers in psychology, 8, 1118.

Marlin, T., Snekser, J. L., & Leese, J. M. (2019). Juvenile convict cichlids shoaling decisions in relation to shoal size and age. Ethology, 125(7), 485–491.

Maury, O. (2017). Can schooling regulate marine populations and ecosystems?. Progress in Oceanography, 156, 91–103.

Mousseau, T. A., & Roff, D. A. (1987). Natural selection and the heritability of fitness components. Heredity, 59(2), 181–197.

Ogale, S. N. (2002). Mahseer breeding and conservation and possibilities of commercial culture. The Indian experience. FAO Fisheries Technical Paper, 193–212.

Paijmans, K. C., Booth, D. J., & Wong, M. Y. (2019). Towards an ultimate explanation for mixed[species shoaling. Fish and Fisheries, 20(5), 921–933.

Piffer, L., Petrazzini, M. E. M., & Agrillo, C. (2013). Large number discrimination in newborn fish. PLoS One, 8(4).

Pinder, A. C., Britton, J. R., Harrison, A. J., Nautiyal, P., Bower, S. D., Cooke, S. J., … & Walton, S. (2019). Mahseer (Tor spp.) fishes of the world: status, challenges and opportunities for conservation. Reviews in Fish Biology and Fisheries, 29(2), 417–452.

Prasad, A. D., Venkataramana, G. V., & Thomas, M. (2009). Fish diversity and its conservation in major wetlands of Mysore. Journal of Environmental biology, 30(5), 713.

Pritchard, V. L., Lawrence, J., Butlin, R. K., & Krause, J. (2001). Shoal choice in zebrafish, Danio rerio: the influence of shoal size and activity. Animal Behaviour, 62(6), 1085–1088.

Raghavan, R., Ali, A., Dahanukar, N., & Rosser, A. (2011). Is the Deccan Mahseer, *Tor khudree* (Sykes, 1839)(Pisces: Cyprinidae) fishery in the Western Ghats Hotspot sustainable? A participatory approach to stock assessment. Fisheries Research, 110(1), 29–38.

Rodgers, G. M., Ward, J. R., Askwith, B., & Morrell, L. J. (2011). Balancing the dilution and oddity effects: decisions depend on body size. PLoS One, 6(7), e14819.

Salvanes, A. G., Moberg, O., & Braithwaite, V. A. (2007). Effects of early experience on group behaviour in fish. Animal Behaviour, 74(4), 805–811.

Salvanes, A. G. V., Moberg, O., Ebbesson, L. O., Nilsen, T. O., Jensen, K. H., & Braithwaite, V. A. (2013). Environmental enrichment promotes neural plasticity and cognitive ability in fish. Proceedings of the Royal Society B: Biological Sciences, 280(1767), 20131331.

Seguin, D., & Gerlai, R. (2017). Zebrafish prefer larger to smaller shoals: analysis of quantity estimation in a genetically tractable model organism. Animal cognition, 20(5), 813–821.

Sreekantha, K. V., & Ramachandra, T. V. (2005). Fish diversity in Linganamakki Reservoir, Sharavathi River. Ecol Environ Conserv, 11, 337–348.

Sridhar, H., & Guttal, V. (2018). Friendship across species borders: factors that facilitate and constrain heterospecific sociality. Philosophical Transactions of the Royal Society B: Biological Sciences, 373(1746), 20170014.

Stamps, J. A., & Biro, P. A. (2016). Personality and individual differences in plasticity. Current opinion in behavioral sciences, 12, 18–23.

Stephenson, J. F. (2019). Parasite-induced plasticity in host social behaviour depends on sex and susceptibility. Biology letters, 15(11), 20190557.

Sundström, L. F., Petersson, E., Höjesjö, J., Johnsson, J. I., & Järvi, T. (2004). Hatchery selection promotes boldness in newly hatched brown trout (*Salmo trutta*): implications for dominance. Behavioral Ecology, 15(2), 192–198.

Thambithurai, D., Hollins, J., Van Leeuwen, T., Rácz, A., Lindström, J., Parsons, K., & Killen, S. S. (2018). Shoal size as a key determinant of vulnerability to capture under a simulated fishery scenario. Ecology and evolution, 8(13), 6505–6514.

Toms, C. N., Echevarria, D. J., & Jouandot, D. J. (2010). A methodological review of personality-related studies in fish: focus on the shy-bold axis of behavior. International Journal of Comparative Psychology, 23(1), 1–25.

Ullah, I., Zuberi, A., Khan, K. U., Ahmad, S., Thörnqvist, P. O., & Winberg, S. (2017). Effects of enrichment on the development of behaviour in an endangered fish mahseer (*Tor putitora*). Applied animal behaviour science, 186, 93–100.

Varma, V., Vasoya, H., Jain, A., & Binoy, V. V. (2020a, *In Press*). ‘The Bold are the Sociable’: Personality and lateralized utilization of brain hemisphere in the juveniles of a megafish Deccan Mahseer (*Tor khudree*). Ichthyological Research, https://doi.org/10.1007/s10228-020-00744-8.

Varma, V., Vasoya, H., Jain, A., & Binoy, V. V. (2020). Big fish in small tanks: Stunting in the Deccan Mahseer, *Tor khudree* (Sykes 1849). bioRxiv, https://doi.org/10.1101/2020.04.04.025049

Vila Pouca, C., & Brown, C. (2019). Lack of social preference between unfamiliar and familiar juvenile Port Jackson sharks *Heterodontus portusjacksoni*. Journal of fish biology, 95(2), 520–526.

Ward, A. J., Kent, M. I., & Webster, M. M. (2020). Social recognition and social attraction in group-living fishes. Frontiers in Ecology and Evolution, 8, 15.

Way, G. P., Kiesel, A. L., Ruhl, N., Snekser, J. L., & McRobert, S. P. (2015). Sex differences in a shoaling-boldness behavioral syndrome, but no link with aggression. Behavioural processes, 113, 7–12.

Webster, M. M., Goldsmith, J., Ward, A. J. W., & Hart, P. J. B. (2007). Habitat-specific chemical cues influence association preferences and shoal cohesion in fish. Behavioral Ecology and Sociobiology, 62(2), 273–280.

Yoshida, M., Nagamine, M., & Uematsu, K. (2005). Comparison of behavioral responses to a novel environment between three teleosts, bluegill *Lepomis macrochirus*, crucian carp *Carassius langsdorfii*, and goldfish *Carassius auratus*. Fisheries Science, 71(2), 314–319.

